# A lipid-based parallel processor for chemical signals

**DOI:** 10.1101/2021.05.05.442835

**Authors:** Idil Cazimoglu, Michael J. Booth, Hagan Bayley

## Abstract

A key goal of bottom-up synthetic biology is to construct cell- and tissue-like structures. Underpinning cellular life is the ability to process several external chemical signals, often in parallel. Until now, however, cell- and tissue-like structures have only been constructed with one signalling pathway. Here, we construct a dual-signal processor from the bottom up in a modular fashion. The processor comprises three aqueous compartments bounded by lipid bilayers and operates in an aqueous environment. It can receive two chemical signals from the external environment, process them orthogonally, and then produce a distinct output for each signal. It is suitable for both sensing and enzymatic processing of environmental signals with fluorescence and molecular outputs. In the future, such processors could serve as smart drug delivery vehicles or as modules within synthetic tissues to control their behaviour in response to external chemical signals.

## Introduction

Lipid-bounded aqueous compartments have applications in drug delivery as well as bottom-up synthetic biology. Single compartment structures are interfaced to an external aqueous environment through a lipid bilayer and may contain sub-compartments^1^. Those devised for drug delivery are designed to release their contents. The release can be passive through biodegradation^2^ or targeted through structural degradation coupled to environmental triggers such as pH, ultrasound or other external signals^2–5^. More sophisticated single compartments exhibit cell-like features. For example, they can employ pore-forming membrane proteins to release contents without structural degradation in response to a trigger^1,5^. They may also receive signals from the environment to activate internal chemical processes such as ATP generation^6,7^, protein expression through transcription and translation^7,8^ or glucose metabolism^9^.

Multi-compartment structures might execute more complex functions, by acting as synthetic tissues^10^. Additional compartments not only can carry out an increased number of individual functions, but also can collectively exhibit emergent properties^11,12^. Much of the work on multi-compartment structures has been conducted in an external oil environment. When two aqueous compartments are placed within a lipid-containing oil, lipid monolayers form around each of them. Hence, when two such compartments are brought together a lipid bilayer forms between them, termed droplet interface bilayer (DIB)^13^. Networks of droplets interconnected by DIBs can be generated manually^14,15^ or by 3D-printing^12,16^. Within oil, these structures can process external light^15,17^, mechanical^18^ or electrical^11^ stimuli, but they cannot handle input from water-soluble signalling molecules.

To function under physiological conditions, for example as smart drug delivery vehicles, multi-compartment structures must operate in an aqueous environment and respond to external chemical signals. Accordingly, multisomes, structures consisting of aqueous compartments inside a lipid-containing oil drop, have been generated in water^19,20^. One way of forming them is suspending a lipid-containing oil drop on a Teflon-coated silver wire loop and placing the internal compartments inside the oil drop^19,21^. Multisomes have been shown to degrade by design in response to an external pH or temperature change^19^, and to receive^19^ or send^21^ chemical signals from and to the external environment through pores formed by alpha-hemolysin (αHL) in their lipid bilayers. Multi-compartment vesicles, structures composed of compartments interfaced with one another and the external aqueous environment by lipid bilayers without a surrounding oil drop, have been shown to receive a single chemical signal from the environment, through αHL pores, and activate a single cascade reaction^22^. Therefore, these structures only contained a single signalling pathway and carried out a single task. Natural cells and tissues receive, process and produce several chemical signals, often simultaneously, with little or no cross-talk^23–25^. However, synthetic structures that can simultaneously or independently process multiple external signals to carry out multiple tasks have not been generated.

Here, we produce multi-compartment processors with dedicated signal transmission and processing compartments (**Fig. 1a, f**). We construct these structures from the bottom up in a modular fashion and show that they can receive and process external chemical signals separately or simultaneously and then produce distinct outputs. The output can be fluorescence, suitable for sensing applications, or the production and release of a molecule, suitable for drug delivery applications or chemical communication. Our processors employ the multisome platform (**Fig. 1**). When the three aqueous compartments sink inside the oil drop, interface bilayers form between them as well as between each compartment and the external aqueous environment. The signal transmission compartment contains αHL, which forms pores connecting this compartment to the external solution and the two processing compartments. External pores allow the exchange of chemicals between the environment and the signal transmission compartment, whereas the internal pores allow the exchange of chemicals between the signal transmission compartment and the processing compartments. The functionality of the three-compartment processor is reminiscent of a dual-core central processing unit (CPU) with a bus interface and two processing units, where the bus interface handles the flow of information between the rest of the computer and the CPU cores through the front and back side buses^26^ (**Fig. 1b**). Chemical input signals 1 and/or 2 introduced from the external aqueous environment diffuse through the external and internal pores and reach both processing compartments (**Fig. 1c-e**). Input signal 1 produces a fluorescence output in the sensing compartment (**Fig. 1c**). Input signal 2 is converted by the enzymatic processing compartment and the molecular output diffuses through the internal and external pores into the external environment (**Fig. 1d**). When both input signals are introduced, both outputs are produced simultaneously (**Fig. 1e**).

**Fig. 1:**
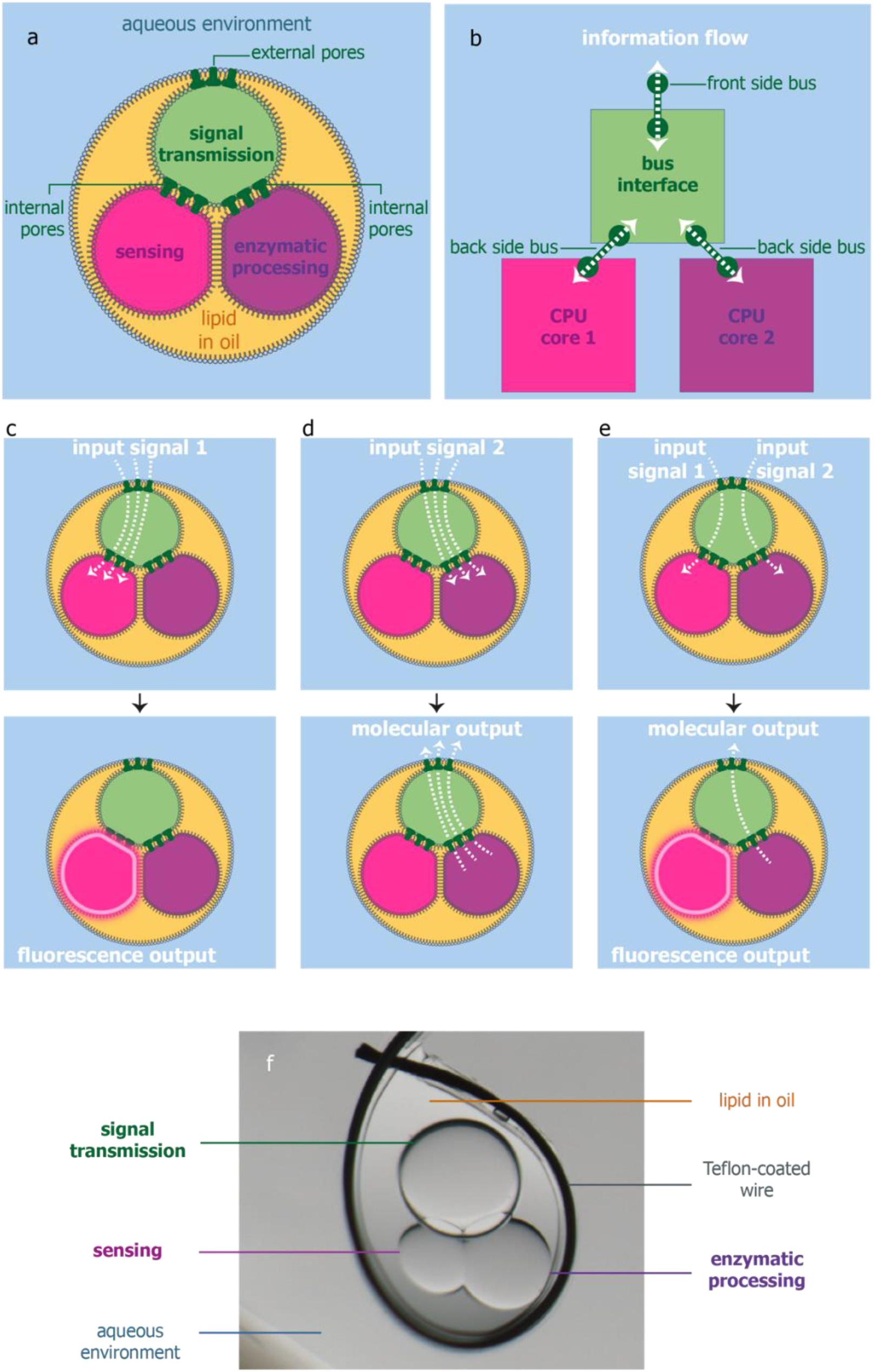
Three-compartment processor with sensing and enzymatic processing compartments. **a** Three-compartment processor for chemical signals containing sensing and enzymatic processing compartments. A signal transmission compartment enables communication between the processing compartments and the external environment. **b** Diagram of a dual-core CPU, analogous to the three-compartment processor. **c** Sensing in a three-compartment processor by intake of input signal 1 and production of a fluorescence output in the sensing compartment. **d** Enzymatic processing in a three-compartment processor by intake of input signal 2 and production of a molecular output and its release into the external environment by the enzymatic processing compartment. **e** Simultaneous sensing and enzymatic processing in a three-compartment processor by intake of both input signals and production of two distinct outputs. **f** Top view of a three-compartment processor contained in an oil drop suspended on a Teflon-coated silver wire loop within an aqueous environment. Wire diameter = 76 μm.

## Results

### Fast molecular diffusion through pores between compartments

The most crucial requirement for our chemical signal processors is effective signal transmission. Each chemical input signal must diffuse through two bilayers to reach a processing compartment. After enzymatic processing, the molecular output must then diffuse through two bilayers again to reach the external environment (a total of 4 bilayers), where it becomes diluted by ~20,000-fold before detection. These factors make fast molecular diffusion essential, which in turn requires efficient insertion of pores into the bilayers.

Diffusion through αHL pores has been shown with Ca^2+^ ions^1,19,27^ and a range of small molecules^21,28–30^. In these cases, the pores were produced after cell-free expression of αHL monomers, by using heptamers from *S. aureus* purified by a lengthy procedure, or by using up to 50-60 μg mL^−1^ of commercially sourced monomers from *S. aureus*. Incomplete diffusion across lipid bilayers was observed in tens of minutes to hours or days. To make our processors feasible, we required much faster diffusion rates.

We expressed recombinant αHL in *E. coli* and separated the monomers and heptamers by size exclusion chromatography. We studied diffusion of 2-[*N*-(7-nitrobenz-2-oxa-1,3-diazol-4-yl) amino]-2-deoxy-D-glucose (2-NBDG) molecules across droplet interface bilayers within an oil external environment containing 1,2-diphytanoyl-sn-glycero-3-phosphatidylcholine (DPhPC) (**Fig. 2a**). Mimicking the dimensions of a processor, we formed 250–300 μm diameter signal release compartments containing 1 mM 2-NBDG and 500–650 μm diameter signal transmission compartments containing 200 μg mL^−1^ αHL monomers (**Fig. 2b**) or 200 μg mL^−1^ heptamers (**Fig. 2c**) or no αHL (**Fig. 2d**). Using purified αHL monomers (**Fig. 2b**), we observed molecular diffusion at unprecedented speeds, with complete equilibration of 2-NBDG molecules within 6 minutes (*n* = 5). We also observed that 2-NBDG diffusion began immediately upon contact of the two compartments and proceeded simultaneously with bilayer formation, as indicated by the increasing contact angle between the compartments^16^ (**Supplementary Fig. 1, Supplementary Video 1**). With αHL heptamers (*n* = 3, **Fig. 2c**) or no αHL (*n* = 3, **Fig. 2d**), transfer of 2-NBDG was not visible after 3 days. The high concentration of αHL, 200 μg mL^−1^, inside the compartments had no adverse effects on the structures or those shown later in this work.

**Fig. 2:**
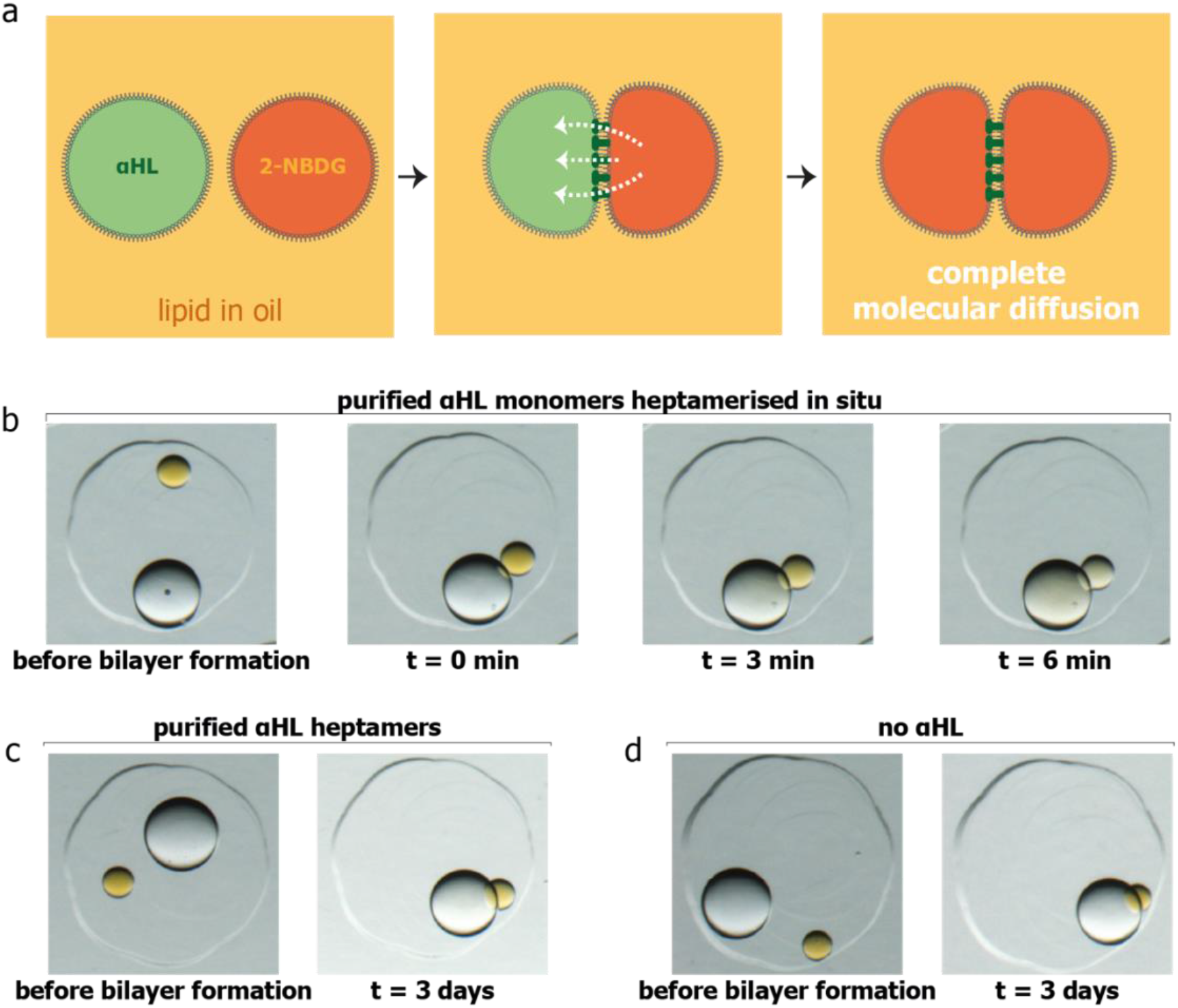
Fast diffusion of molecular signals across lipid bilayers. **a** Bilayer formation between a signal transmission and a signal release compartment within a lipid-oil external environment, leading to diffusion of 2-NBDG molecules through α-hemolysin (αHL) pores. **b-d** Bright-field microscopy images of molecular diffusion from compartments containing 2-NBDG into signal transmission compartments containing purified αHL monomers (**b**), purified αHL heptamers (**c**), and no αHL (**d**).

### Exchange of chemical signals with the external environment

After establishing internal signal transmission between compartments in a lipid-oil external environment, we constructed structures within an aqueous environment. Any chemical output generated by these structures must diffuse through two lipid bilayers, first into the signal transmission compartment, and then into the external aqueous environment. To mimic this process, we built two-compartment structures with a signal release compartment containing 2-NBDG and a signal transmission compartment containing 200 μg mL^−1^ purified αHL monomers (**Fig. 3a**) or no αHL. Complete release of 1 mM 2-NBDG through two bilayers to the external environment was observed within 10 minutes (*n* = 3, **Fig. 3b, Supplementary Video 2**). Without αHL monomers in the signal transmission compartment, 2-NBDG remained in its original compartment (*n* = 3, **Fig. 3c**).

**Fig. 3:**
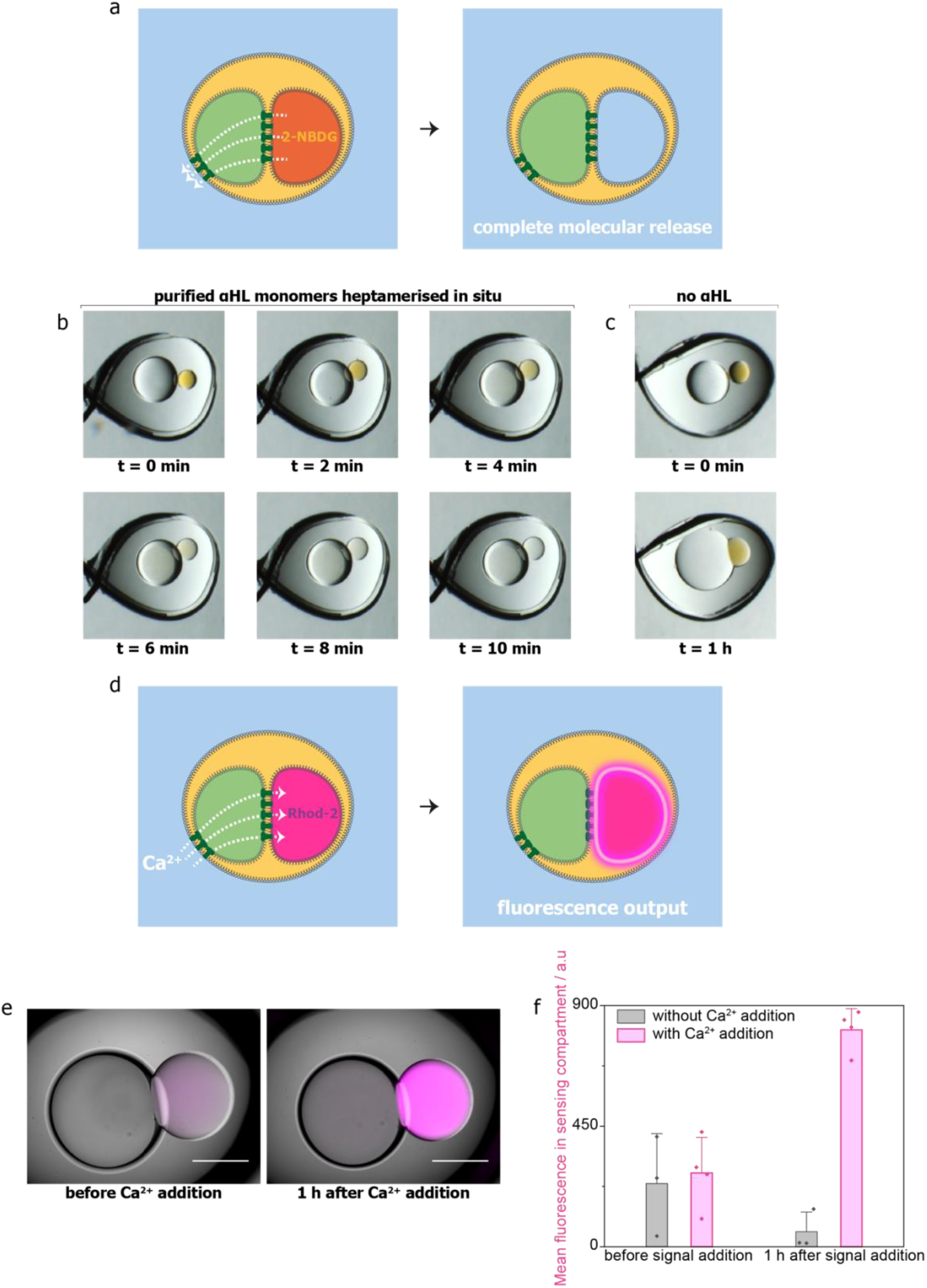
Output and intake of chemical signals in an external aqueous environment. **a** Output of chemical signal 2-NBDG through a signal transmission compartment, leading to its release into the external aqueous environment. **b-c** Bright-field microscopy images of 2-NBDG release into the external aqueous environment through signal transmission compartments with purified αHL monomers (**b**) and without αHL (**c**). Wire diameter = 76 μm. **d** Intake of chemical signal Ca^2+^ through the signal transmission compartment, leading to a fluorescence output in the sensing compartment, which contains dextran-conjugated Rhod-2. **e** Composite bright-field and epifluorescence images of signal transmission from the external environment into the sensing compartment before and 1 h after Ca^2+^ addition to the external aqueous environment. Scale bars = 300 μm. **f** Mean fluorescence values of the sensing compartment before and 1 h after Ca^2+^ addition without (*n* = 3) and with (*n* = 4) Ca^2+^ addition. Error bars represent the standard deviation.

Our processor would also need to receive external signals through two bilayers, first into the signal transmission compartment, then into the sensing compartment to produce a fluorescence output (**Fig. 3d**). To show signal intake and sensing, we built two-compartment structures with a signal transmission compartment containing 200 μg mL^−1^ αHL monomers, and a sensing compartment containing 20 μM dextran-conjugated Ca^2+^ indicator Rhod-2 (~11,000 Da). To minimise reagent use, we included a smaller (300–350 μm diameter) sensing compartment, compared to the larger (500– 650 μm) signal transmission compartment. Fluorescence of the Rhod-2 in the sensing compartment increased 3-fold 1 h after 10 mM Ca^2+^ addition (**Fig. 3e, f**) (*n* = 4). When no Ca^2+^ input signal was added, the fluorescence in the sensing compartment did not increase (*n* = 3, **Fig. 3f**). In fact, in this negative control, the fluorescence decreased as a result of excess chelator in the external solution, initially added to prevent the binding of trace metal ions to Rhod-2, diffusing into the sensing compartment (see **Supplementary Note**).

### Input-activated enzymatic reaction with fluorescence output

The next step was to couple signal intake with activation of an enzymatic process. We chose a restriction endonuclease, EcoRI, which requires Mg^2+^ as a co-factor^31^, and had not been encapsulated in synthetic biological systems before. As a substrate, we designed a molecular beacon based on a previously published sequence^32^: a DNA hairpin with a fluorophore attached on the 5’ end and a quencher on the 3’ end, containing an EcoRI cleavage site 4 and 8 bases away from the 5’ and 3’ ends, respectively^32^. To produce high quenching efficiency^33^, we used the fluorophore Cyanine 5 and the quencher BHQ-3^34^. Upon cleavage at the EcoRI site, the fluorophore/quencher pair separates due to DNA denaturation, resulting in a fluorescence signal (**Fig. 4a**).

**Fig. 4:**
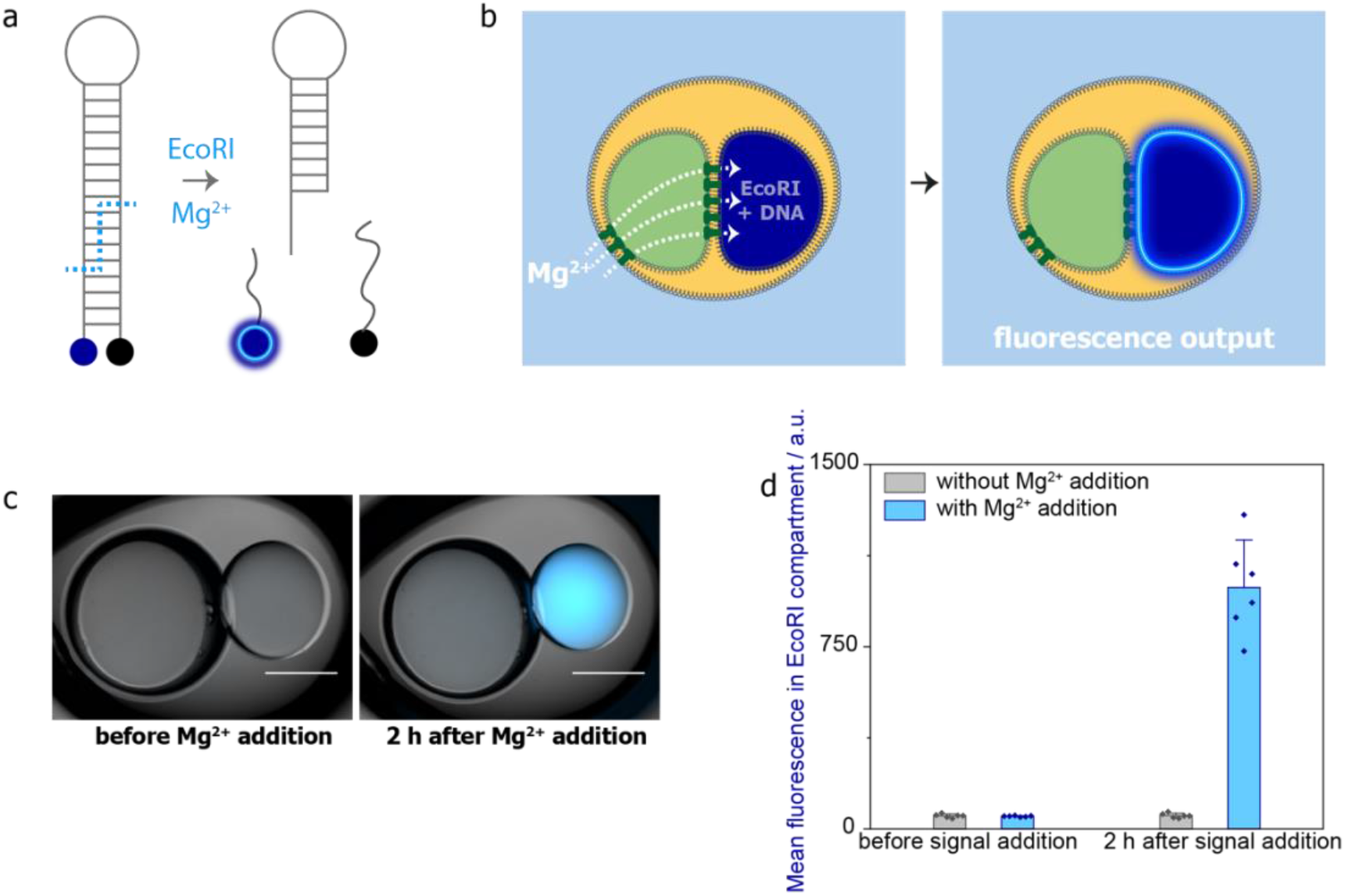
Enzymatic processing by EcoRI in two-compartment processors: signal intake, signal processing and fluorescence output. **a** Design of a Mg^2+^-dependent enzymatic process with a fluorescence output. Upon addition of the input signal, a molecular beacon is cleaved by the endonuclease EcoRI, leading to fluorophore/quencher pair separation and a fluorescence output signal. **b** Intake of ionic signal Mg^2+^ from the external aqueous environment into a two-compartment processor, and activation of enzymatic processing by EcoRI to produce a fluorescence output. **c** Composite bright-field and epifluorescence images of a two-compartment processor containing EcoRI and the DNA substrate before and after Mg^2+^ addition into the external aqueous environment and 2 h at 37 °C. Scale bars = 300 μm. **d** Mean fluorescence values of the EcoRI-containing compartment before and 2 h after Mg^2+^ addition and incubation at 37 °C without (*n* = 6) and with (*n* = 6) Mg^2+^ addition. Error bars represent the standard deviation.

We built two-compartment processors with a signal transmission compartment containing αHL monomers, and an enzymatic processing compartment containing 400 U mL^−1^ EcoRI and 400 nM DNA substrate. To minimise reagent use, we included 300–350 μm diameter enzymatic processing compartments and 500–650 μm diameter signal transmission compartments containing 200 μg mL^−1^ αHL monomers. Mg^2+^ input signal added to the external aqueous environment diffused through the signal transmission compartment into the processing compartment where it induced the cleavage reaction and the fluorescence output (**Fig. 4b**). Fluorescence in the EcoRI compartment increased 19-fold 2 h after 10 mM Mg^2+^ addition (**Fig. 4c, d**) and incubation at 37 °C (*n* = 6). When no Mg^2+^ was added, the fluorescence in the EcoRI compartment remained constant (*n* = 6**, Fig. 4d**). Notably, we obtained stable structures despite the use of 100 μg mL^−1^ bovine serum albumin (BSA) in the enzymatic processing compartment. For these structures, we initially used 200 μg mL^−1^ αHL monomers in the signal transmission compartment and did not observe an increase in fluorescence in the EcoRI compartment after the addition of external Mg^2+^. We made new structures with a reduced αHL concentration (see Methods) and were then able to detect the fluorescence signal (**Fig. 4c, d**). This indicated that the fluorescent product was diffusing out of the reaction compartment through the pores, which we confirmed by fluorescence measurements on the signal transmission compartment (**Supplementary Fig. 2**).

### Input-activated enzymatic reaction with molecular output

Having shown ionic signal intake and enzymatic activation in two-compartment processors, we next aimed to demonstrate molecular signal intake, enzymatic turnover, and output of the product molecule into the external environment. We built two-compartment processors with a signal transmission compartment containing 200 μg mL^−1^ αHL monomers, and an enzymatic processing compartment containing 700 μg mL^−1^ (30 U mL^−1^) β-galactosidase, which hydrolyses lactose into glucose and galactose (**Fig. 5a**). The lactose input signal, added to the external aqueous environment, diffused through two bilayers, first to the signal transmission compartment and then to the processing compartment where it was hydrolysed. The product glucose was then released back through the same two bilayers into the external environment as a molecular output (**Fig. 5b**). In our set up, the external environment was 800 μL in volume. As the product glucose would be diluted upon release to the external environment, we included larger enzymatic processing compartments (400–470 μm in diameter, ~40 nL in volume) than those we previously used for fluorescence output. Using these larger β-galactosidase compartments, the product glucose was still diluted by ~20,000-fold upon release. We used signal transmission compartments of the same size as previously used. After 40 mM lactose addition and incubation at 37 °C for 6 h, glucose was detected in the external environment by using a commercial assay (*n* = 6, **Fig. 5c**). When no lactose was added, no glucose was detected (*n* = 6, **Fig. 5c**). In processors incubated with lactose, the β-galactosidase compartment shrank (**Fig. 5d**). We attribute this observation to the increased osmotic pressure of the external aqueous environment. On the other hand, the signal transmission compartment, interfaced directly with the external environment through αHL pores, was better able to take in solutes and reach osmotic equilibrium. The processors remained intact despite the volume changes.

**Fig. 5:**
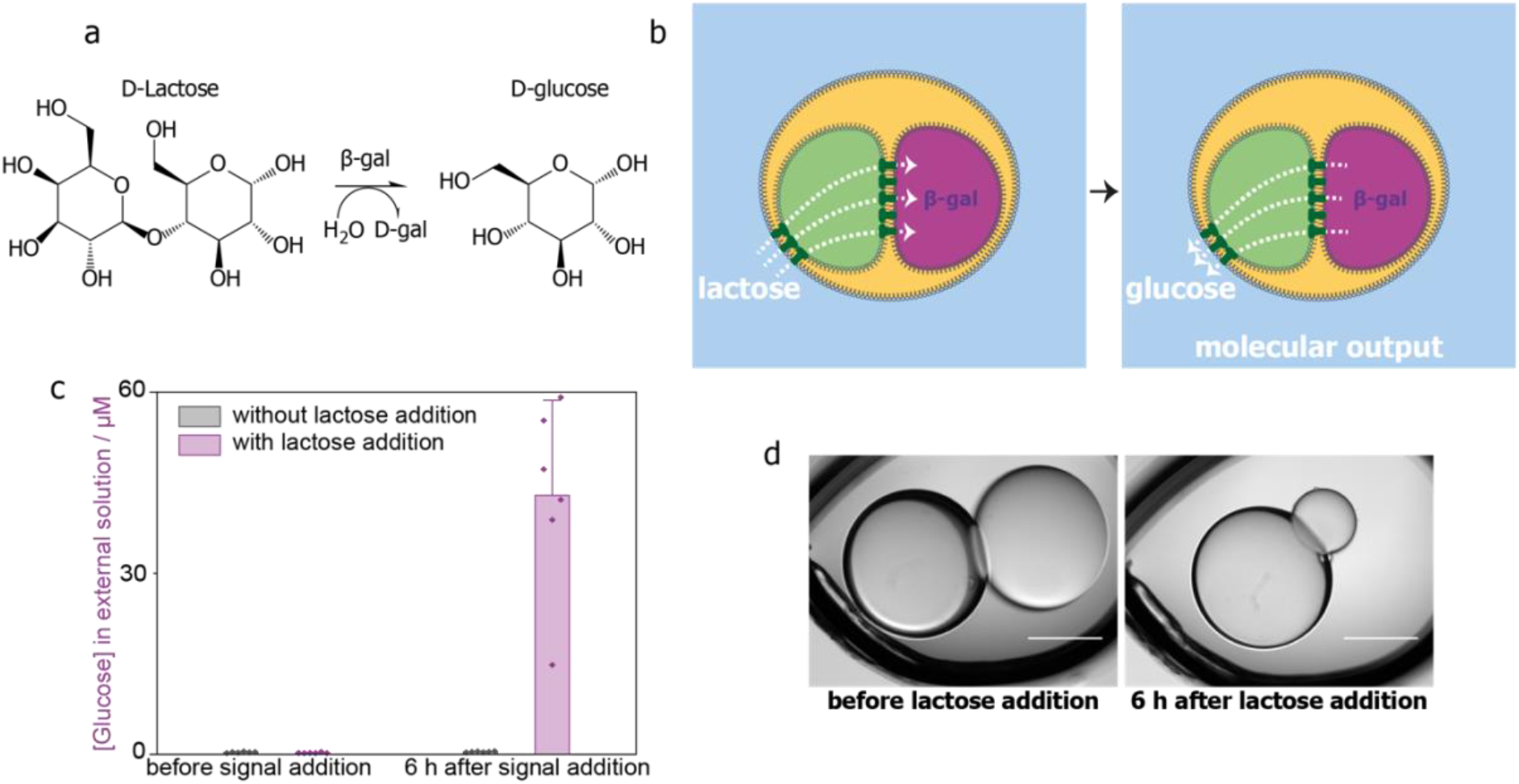
Enzymatic processing by β-galactosidase in two-compartment processors: signal intake, signal processing, small molecule production and release. **a** Hydrolysis of D-lactose by β-galactosidase (β-gal) produces D-galactose (D-gal) and D-glucose. **b** Intake of the molecular input signal lactose from the external environment into a two-compartment processor, its enzymatic processing and release of the product glucose into the external environment. **c** Concentration of glucose in the external aqueous environment before and 6 h after lactose addition and incubation at 37 °C, without (*n* = 6) and with (*n* = 6) lactose addition. Error bars represent the standard deviation. **d** Bright-field microscopy images of a two-compartment processor containing β-galactosidase before and after lactose addition and processing. Scale bars = 300 μm.

### Orthogonal processing of two input signals

At this stage, we had all the parts required to build a processor that would independently and simultaneously process two external signals. Combining a signal transmission compartment containing 200 μg mL^−1^ αHL monomers, a sensing compartment containing 20 μM dextran-conjugated Rhod-2, and an enzymatic processing compartment containing 700 μg mL^−1^ β-galactosidase, we constructed three-compartment processors (**Fig. 6a**). We tested these processors with all four possible input signal combinations: no input signals, only Ca^2+^, only lactose, and both Ca^2+^ and lactose (**Fig. 6b**). When no input signals were added, fluorescence in the Rhod-2 compartment decreased and no glucose was detected in the external environment, after 3 h at 37 °C (*n* = 4). Addition of only the Ca^2+^ input signal led to a fluorescence output in the Rhod-2 compartment and no glucose was detected in the external environment (*n* = 5). When only the lactose input signal was added, the fluorescence of the Rhod-2 compartment decreased, and glucose was detected in the external environment (*n* = 4). When both input signals were added, fluorescence in the Rhod-2 compartment increased (**Fig. 6b, c**) and glucose was detected in the external environment (*n* = 5, **Fig. 6b**). These results demonstrate the orthogonal processing of two chemical inputs within a lipid-bound multicompartment structure.

**Fig. 6:**
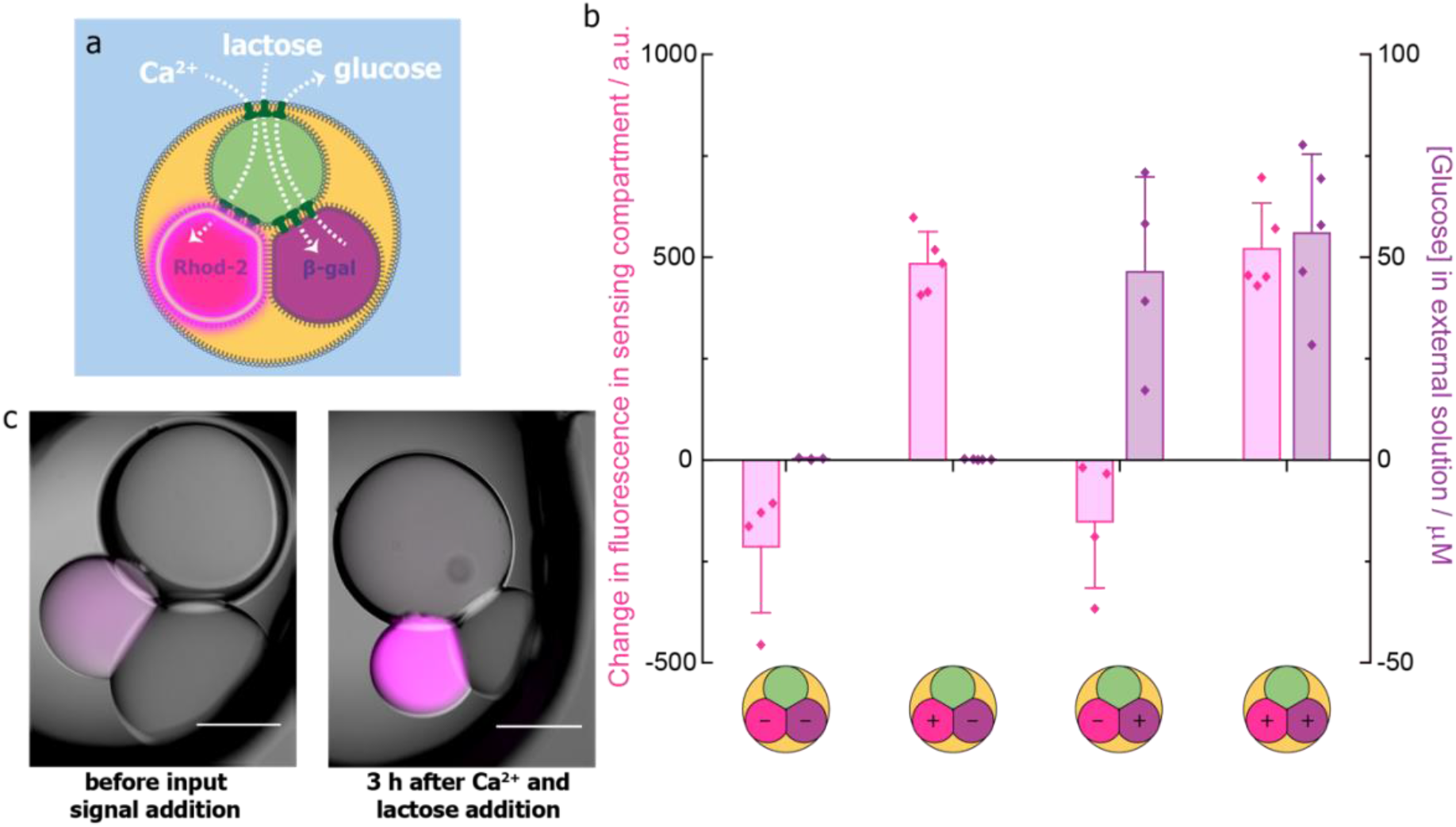
Independent and simultaneous sensing and enzymatic processing in three-compartment processors. **a** Three-compartment processor containing signal transmission, sensing and enzymatic processing compartments. The intake of Ca^2+^ leads to a fluorescence output in the sensing compartment containing Rhod-2, whereas the intake of lactose leads to the production of glucose by β-galactosidase and its release into the external environment. **b** Changes in the mean fluorescence of Rhod-2 compartments and the mean concentration of glucose detected in the external solution after 3 h at 37 °C for three-compartment processors with no input signals (*n* = 4) and with an input of Ca^2+^ only (*n* = 5), lactose only (*n* = 4), and both Ca^2+^ and lactose (*n* = 5). Error bars represent the standard deviation. **c** Composite bright-field and epifluorescence images of a three-compartment processor with a sensing compartment containing Rhod-2 and an enzymatic processing compartment containing β-galactosidase before and after simultaneous addition of Ca^2+^ and lactose input signals and 3 h at 37 °C. Scale bars = 300 μm.

### Simultaneous enzymatic processing of two input signals

To demonstrate simultaneous enzymatic processing, we constructed processors with a signal transmission compartment containing 200 μg mL^−1^ αHL monomers and two enzymatic processing compartments: one containing 400 U mL^−1^ EcoRI with 400 nM DNA substrate and one containing 700 μg mL^−1^ β-galactosidase (**Fig. 7a**). We had to find a pH value that would allow both reactions to proceed, and selected pH 6.5 as a compromise (**Supplementary Fig. 3, 4**). The addition of both input signals, 10 mM Mg^2+^ and 40 mM lactose, followed by incubation at 37 °C for 3 h generated increased fluorescence in the EcoRI compartment (**Fig. 7b, c)** and glucose in the external environment (**Fig. 7b**) simultaneously (*n* = 3), despite the suboptimal reaction conditions.

**Fig. 7:**
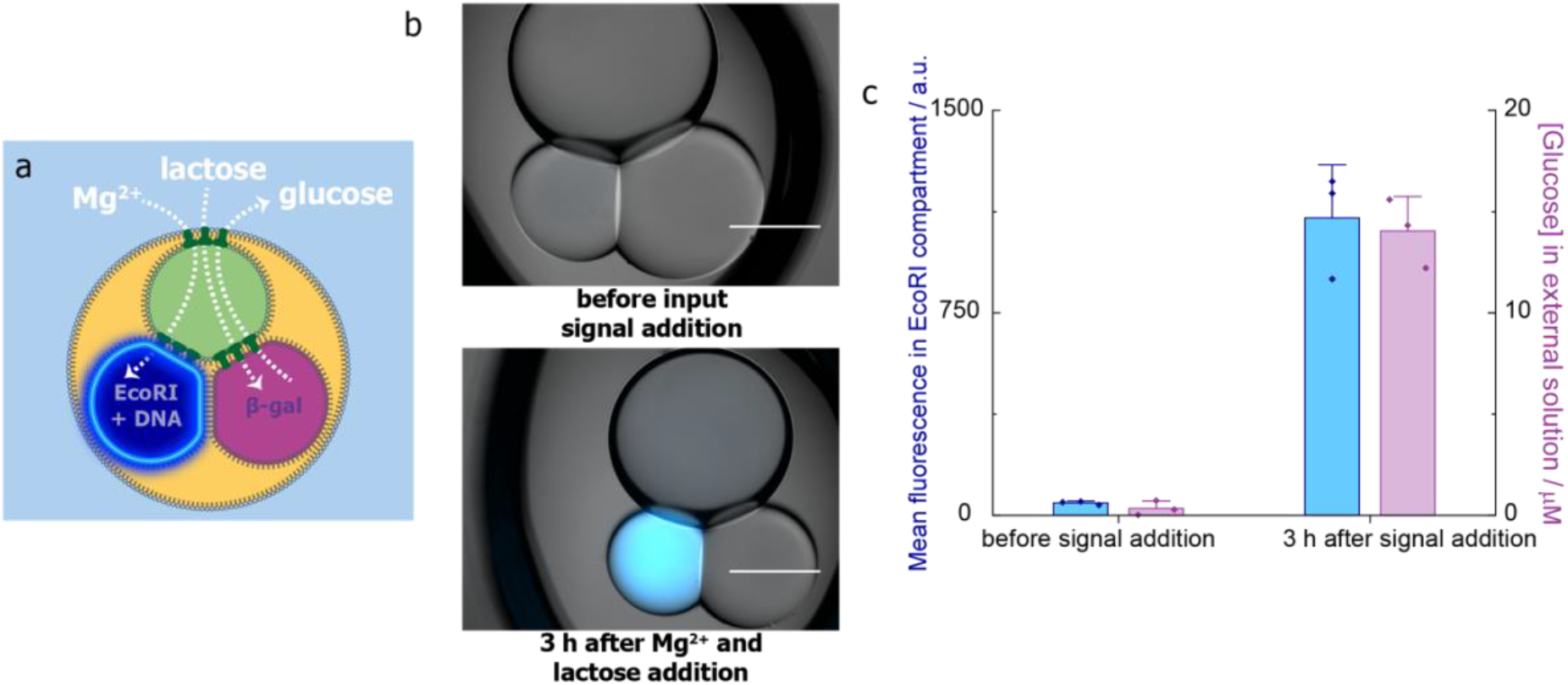
Simultaneous enzymatic processing in three compartment processors. **a** Three-compartment processor for simultaneous enzymatic processing by EcoRI and β-galactosidase activated by input signals Mg^2+^ and lactose, respectively. **b** Composite bright-field and epifluorescence images of a three-compartment processor with EcoRI and β-galactosidase processing compartments before and after the simultaneous addition of Mg^2+^ and lactose and 3 h incubation at 37 °C. Scale bars = 300 μm. **c** Mean fluorescence values in EcoRI-containing compartment and the mean concentration of glucose detected in the external solution before and after the simultaneous addition of Mg^2+^ and lactose and 3 h at 37 °C (*n* = 3).

## Discussion

In summary, we have constructed processors from lipid bilayer-bounded compartments from the bottom up, in a modular fashion. By using purified recombinant αΗL monomers, we achieved fast diffusion of ionic and molecular signals across as many as 4 lipid bilayers. We first demonstrated the release and intake of chemical signals in two-compartment structures. We then encapsulated enzymatic processes within them and with an ionic input signal, we activated DNA cleavage. We also demonstrated the enzymatic hydrolysis of a molecular input signal and release of the product into the external environment as a molecular output. By combining different components in a modular fashion, we built three-compartment processors that receive and process two different chemical signals in an orthogonal manner, producing two distinct outputs: fluorescence and molecule release. We also showed simultaneous activation of different enzymes in two separate compartments of a three-compartment processor. This work represents the first bottom-up synthetic biological system with independent and simultaneous processing of more than one signal, and serves as a steppingstone in the development of multi-compartment systems with complex signal processing capabilities. As a result of the use of the recombinant αΗL monomers introduced in this work, engineered αΗL pores might be used at high concentrations to modulate signal transmission using small molecules^21,35,36^, light^37^ or other stimuli^38,39^, adding further complexity to the tasks performed by future structures.

Our processors are modular, robust and versatile. Lipid-bounded compartments in an external aqueous environment often suffer from structural instability^21,27^, which can be addressed by balancing osmotic pressures during construction^21,29^ but limiting the applicability of these structures in dynamic environments. Our processors, functionalised with fast signal processing capabilities involving enzymatic reactions, can withstand changing osmotic pressures. They also incorporate high concentrations of enzymes and membrane proteins and are stable at 37 °C.

Due to their modularity and robustness, the processors might be adapted for a variety of applications. To simultaneously process complex sets of input signals, several processing units could be connected to a large signal transmission compartment (**Supplementary Fig. 5, Supplementary Video 3**). The platform could be used to build sensors and micro-reactors. For example, the encapsulation of suitable reporters would allow parallel processing in medical diagnostics, water quality analysis, or the sensing of bacteria. Micro-reactors could be built by combining compartments that encapsulate multiple independent reactions, cooperative reactions forming a cascade, or both. The multi-compartment processors might also be incorporated into drug delivery systems or synthetic tissues to enable complex communication with their environment through parallel signal processing. For example, the processors could be engineered to monitor multiple biomarkers and integrate outputs for sophisticated drug delivery. In the future, these processors could also be used to handle signals between interfaced synthetic and living tissues, coordinating them to act as functional hybrid systems.

## Methods

### αHL expression and purification

BL21(DE3)pLysS *E. coli* cells (Agilent) were transformed with ~2 μg αHL-D8H6 plasmid^40^ without heat shock and incubated on LB-agar plates containing 25 g L^−1^ lysogeny broth (LB) and 15 g L^−1^ agar with antibiotics (50 μg mL^−1^ carbenicillin disodium and 34 μg mL^−1^ chloramphenicol) at 37 °C for 16 h. Single colonies were picked and inoculated into 15 g L^−1^ LB with antibiotics at 37 °C with shaking at 250 rpm until OD ≈ 0.7. The cultures were then induced with 1 mM isopropyl β-d-1-thiogalactopyranoside (IPTG) and incubated at 18 °C with shaking at 250 rpm for 16 h. Between this point and Fast Protein Liquid Chromatography (FPLC) purification, all steps were carried out at 4 °C. The cells were pelleted and lysed in 50 mM Tris-HCl, pH 8.0, 0.5 M NaCl, 10 mM imidazole and 0.1% Triton X-100 followed by the addition of ice-cold MgCl_2_ (final concentration, 5 mM), hen egg white lysozyme (final concentration, 1 mg mL^−1^) and 250 units benzonase.

The resulting lysate was sonicated using an ultrasonic probe with 30 s pulses and 30 s intervals for 3 min followed by centrifugation to separate the supernatant and cell debris. The supernatant was loaded onto a column pre-loaded with 2 mL Ni-NTA resin (HisPur) and placed on a rotator disk for 1 h. The column was then washed twice with 15 mL of 50 mM Tris-HCl pH 8.0, 0.5 M NaCl, 10 mM imidazole and 0.1% Triton X-100. Proteins were eluted in 5 × 1 mL batches with 50 mM Tris-HCl pH 8.0, 0.5 M NaCl, 250 mM imidazole and 0.1% Triton X-100 and stored at –80 °C until FPLC purification.

To separate monomers and heptamers and exchange the buffer, 500 μL of an elution with high protein concentration, as judged by SDS-PAGE electrophoresis, was loaded at room temperature onto a Superdex 75 100/300 GL (GE Healthcare) gel filtration column which was equilibrated and run in 10 mM Tris-HCl pH 8.0, 200 mM NaCl at a flow rate of 0.5 mL min^−1^ and eluted as 0.5 mL volume per fraction. Monomeric and heptameric αHL were detected by ultraviolet absorption at 254 and 280 nm, and by SDS-PAGE electrophoresis. Protein concentrations were measured using a Nanodrop. The fractions with highest concentrations (200–400 μg mL^−1^) were stored as 5–10 μL aliquots in Protein LoBind tubes (Eppendorf) at –80 °C.

### Compositions of solutions

1,2-Diphytanoyl-sn-glycero-3-phosphocholine (DPhPC, Avanti Polar Lipids) was stored as powder at – 80 °C. Hexadecane (Merck) and silicone oil AR20 (Wacker) were filtered before use through 0.22 μm polyethersulfone filters (Corning) under vacuum. Lipid-oil solutions were prepared by dissolving the desired amount of lipid in chloroform in isopropanol-cleaned glass vials and evaporating the solvent under a slow stream of nitrogen gas while manually rotating. The films were dried under vacuum overnight and stored under argon in Teflon-capped glass vials (Supelco). For use, a film was dissolved by sonication for 45 min in 65:35 v:v silicone oil:hexadecane to give 2 mM DPhPC (5 mM for Supplementary Fig. 1).

Rhod-2 dextran conjugate (~11,000 Da) was from AAT Bioquest. 2-(*N*-(7-Nitrobenz-2-oxa-1,3-diazol-4-yl)amino)-2-deoxyglucose (2-NBDG), the tetrapotassium salt of BAPTA and the Amplex Red Glucose/Glucose Oxidase Assay Kit were from Invitrogen. EcoRI-HF was from New England Biolabs. All other reagents used in aqueous solutions were purchased from Merck. β-galactosidase from *A. oryzae* was purified with a PD-10 Desalting Column (GE Healthcare). The custom-made DNA substrate was purchased from ATDBio. It was designed based on a sequence described previously^32^, but with the fluorophore Cyanine 5 on the 5’ end and the quencher Black Hole Quencher (BHQ-3) on the 3’ end.

An individual aliquot of αHL was thawed on ice for each experiment and diluted with 10 mM Tris-HCl pH 8.0, 200 mM NaCl to give 200 μg mL^−1^ αHL. For processors involving a compartment containing EcoRI and DNA, the αHL was diluted further to a concentration at which the equilibration of 2-NBDG across a bilayer (Fig. 2) took 20–30 min.

All other compartments and external aqueous environments contained 50 mM MES, 100 mM NaCl at pH 6.5 for experiments with EcoRI or 2-NBDG and pH 5.5 for all other experiments. All processing compartments and external environments contained 2 μM BAPTA for experiments using Rhod-2. NBDG was used at 1 mM. Rhod-2 was used at 20 μM. Compartments containing EcoRI and DNA contained 400 U mL^−1^ EcoRI, 400 nM DNA substrate, 100 μg mL^−1^ BSA. β-galactosidase was used at 700 μg mL^−1^. Input signals were at final concentrations of 0 or 10 mM CaCl_2_ (for Rhod-2), 0 or 10 mM MgCl_2_ (for EcoRI), and 0 or 40 mM D-lactose (for β-galactosidase) in the same buffer conditions as each external solution into which they were added.

### Formation of processors and processor mimics

Chambers were formed as previously described^21^. After the addition of external aqueous solution to each chamber, lipid-containing oil drops (~1.5 μL) were formed on Teflon-coated silver wire loops, zapped with an anti-static gun, and incubated to form a monolayer for at least 5 min. ~0.5 μL of each oil drop was pipetted out to reduce the drop volumes. The oil drops were then incubated for at least another 5 min.

Compartments were formed in the lipid-containing oil inside poly(methyl methacrylate) (PMMA) wells, by using a Gilson P2 pipette. 2-NBDG, β-galactosidase, Rhod-2 compartments were formed first and incubated in the PMMA wells for at least 10 min. EcoRI and αHL compartments were formed later and not incubated. To make each processor, all compartments were simultaneously transferred into the oil drop using a pipette.

For processors with a molecular output, sample solution was pipetted out of each chamber before signal addition for analysis. For all processors, the total external solution after signal addition was 800 μL. The processors were incubated for the specified time, at 37 °C where indicated, then sample solutions were pipetted out of each chamber again for analysis. For all processors, brightfield and or epifluorescence microscopy images were taken before and after signal addition and incubation.

### Fluorescence output detection and analysis

Images for fluorescence outputs were taken using a Leica DMi8 epi fluorescence microscope. For 2-NBDG, the filter cube GFP was used (ex: 450–490 nm, em: 500–550 nm), for Rhod-2 DSRED was used (ex: 540–552, em: 567–643 nm), and for EcoRI+DNA DAPI was used (ex: 325-375, em: 435–485). For quantification, the fluorescence compartments were detected and analysed on Fiji/ImageJ by a custom script followed by manual verification and, in rare cases involving artefacts caused by an air bubble or the silver wire, correction. The script applied Gaussian blur with a standard deviation of 4 and Moments automatic thresholding to each fluorescence image, then used the built-in particle analysis tool of Fiji/ImageJ to detect and record a region of interest (ROI) for each particle with an area of 20000–70000 μm^2^ and a circularity of 0.15–1.00. Each ROI was then applied to its corresponding original fluorescence image to record the average fluorescence value within the compartment area. For two-compartment processors with an EcoRI+DNA compartment, an additional ROI was defined by drawing a line across the diameter of each signal transmission compartment. The ROIs were then applied to their corresponding fluorescence images to obtain fluorescence values across each signal transmission compartment.

### Glucose detection

Glucose detection was performed using the Thermo Fisher Amplex Red Glucose/Glucose Oxidase Assay Kit according to the manufacturer’s instructions. For each sample, a calibration curve with a matching buffer was used. For example, samples from a processor that received a lactose input were compared to a glucose calibration curve that also contained lactose. To avoid the non-linear range of the assay, samples with [glucose] > 30 μM were diluted and re-measured.

### Contact angle measurements

Contact angle measurements were obtained as described previously^16^. Briefly, the contact angle (θ) formed between two compartments was calculated from the radius of each compartment (*R*_1_, *R*_2_), and the centre-to-centre distance (*L*) by using the formula^41^:

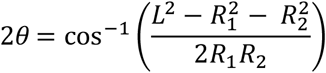

The contact angles were calculated from bright-field microscopy images using a custom-written script in MATLAB (Mathworks), which estimated the radii *R*_1_ and *R*_2_ of the two compartments by circle fitting and the centre-to-centre distance *L* from the distance between the two fitted circles around the compartments. The script was also used to record the average fluorescence value within the detected compartment area.

### EcoRI assays

Using a 96-well plate, 110 μL reaction mixes were prepared in 91 mM NaCl, 91 μg mL^−1^ BSA and 45 mM MES pH 5.5 or MES pH 6.5 or Tris pH 8.0, with 0 or 10 mM MgCl_2_. 364 nM DNA substrate and 364 U mL^−1^ EcoRI were used. The plate was covered and placed in a Tecan Infinite M1000 Pro microplate reader at 37 °C and fluorescence measurements were taken from the bottom of the plate every 5 min with ex = 645–655 nm and em = 665–675 nm.

## Data availability

All relevant data are available from the corresponding authors upon reasonable request.

## Acknowledgements

This research was supported by a European Research Council Advanced Grant. I.C. acknowledges funding by the University of Oxford, the EPSRC & BBSRC Centre for Doctoral Training in Synthetic Biology (grant EP/L016494/1), the Clarendon Fund Scholarship, and the Oxford-Broomhead Graduate Scholarship. M.J.B. acknowledges funding by a Royal Society University Research Fellowship. The authors thank Dr Alessandro Alcinesio for insightful discussions, contact angle analyses and assistance with fluorescence microscopy.

## Author Information

### Affiliations

Department of Chemistry, University of Oxford, Chemistry Research Laboratory, 12 Mansfield Road, Oxford, OX1 3TA, UK

### Contributions

I.C., M.J.B. and H.B. conceived the experiments. I.C. conducted the experiments, collected the data and performed the analyses. I.C., M.J.B. and H.B. wrote the paper.

### Corresponding authors

Correspondence to Hagan Bayley or Michael Booth.

## References

1. Hindley, J. W. et al. Building a synthetic mechanosensitive signaling pathway in compartmentalized artificial cells. Proc. Natl Acad. Sci. USA 116, 16711–16716 (2019).

2. Bulbake, U., Doppalapudi, S., Kommineni, N. & Khan, W. Liposomal Formulations in Clinical Use: An Updated Review. Pharmaceutics 9, 12 (2017).

3. Lee, Y. & Thompson, D. H. Stimuli-responsive liposomes for drug delivery. Wiley Interdiscip. Rev. Nanomed. Nanobiotechnol. 9, e1450 (2017).

4. Rideau, E., Dimova, R., Schwille, P., Wurm, F. R. & Landfester, K. Liposomes and polymersomes: a comparative review towards cell mimicking. Chem. Soc. Rev. 47, 8572–8610 (2018).

5. Haylock, S. et al. Membrane protein mediated bilayer communication in networks of droplet interface bilayers. Commun. Chem. 3, 1–8 (2020).

6. Lee, K. Y. et al. Photosynthetic artificial organelles sustain and control ATP-dependent reactions in a protocellular system. Nat. Biotechnol. 36, 530–535 (2018).

7. Berhanu, S., Ueda, T. & Kuruma, Y. Artificial photosynthetic cell producing energy for protein synthesis. Nat. Commun. 10, 1–10 (2019).

8. Toparlak, D. et al. Artificial cells drive neural differentiation. Sci. Adv. 6, 4920–4938 (2020).

9. Chen, Z. et al. Synthetic beta cells for fusion-mediated dynamic insulin secretion. Nat. Chem. Biol. 14, 86–93 (2018).

10. Bayley, H., Cazimoglu, I. & Hoskin, C. E. G. Synthetic tissues. Emerg. Top. Life Sci. 3, 615–622 (2019).

11. Maglia, G. et al. Droplet networks with incorporated protein diodes show collective properties. Nat. Nanotechnol. 4, 437–440 (2009).

12. Villar, G., Graham, A. D. & Bayley, H. A tissue-like printed material. Science 340, 48–52 (2013).

13. Booth, M. J., Restrepo Schild, V., Downs, F. G. & Bayley, H. Functional aqueous droplet networks. Mol. Biosyst. 13, 1658–1691 (2017).

14. Funakoshi, K., Suzuki, H. & Takeuchi, S. Lipid Bilayer Formation by Contacting Monolayers in a Microfluidic Device for Membrane Protein Analysis. Anal. Chem. 78, 8169–8174 (2006).

15. Holden, M. A., Needham, D. & Bayley, H. Functional Bionetworks from Nanoliter Water Droplets. J. Am. Chem. Soc. 129, 8650–8655 (2007).

16. Alcinesio, A. et al. Controlled packing and single-droplet resolution of 3D-printed functional synthetic tissues. Nat. Commun. 11, 1–13 (2020).

17. Booth, M. J., Schild, V. R., Graham, A. D., Olof, S. N. & Bayley, H. Light-activated communication in synthetic tissues. Sci. Adv. 2, e1600056 (2016).

18. Tamaddoni, N., Freeman, E. C. & Sarles, S. A. Sensitivity and directionality of lipid bilayer mechanotransduction studied using a revised, highly durable membrane-based hair cell sensor. Smart Mater. Struct. 24, 065014 (2015).

19. Villar, G., Heron, A. J. & Bayley, H. Formation of droplet networks that function in aqueous environments. Nat. Nanotechnol. 6, 803–808 (2011).

20. Elani, Y., Solvas, X. C. I., Edel, J. B., Law, R. V. & Ces, O. Microfluidic generation of encapsulated droplet interface bilayer networks (multisomes) and their use as cell-like reactors. Chem. Commun. 52, 5961–5964 (2016).

21. Booth, M. J., Cazimoglu, I. & Bayley, H. Controlled deprotection and release of a small molecule from a compartmented synthetic tissue module. Commun. Chem. 2, 1–8 (2019).

22. Elani, Y., Law, R. V. & Ces, O. Vesicle-based artificial cells as chemical microreactors with spatially segregated reaction pathways. Nat. Commun. 5, 5305 (2014).

23. Asthagiri, A. R. & Lauffenburger, D. A. Bioengineering Models of Cell Signaling. Annu. Rev. Biomed. Eng. 2, 31–53 (2000).

24. Kholodenko, B. N. Cell-signalling dynamics in time and space. Nat. Rev. Mol. Cell Biol. 7, 165–176 (2006).

25. Purvis, J. E. & Lahav, G. Encoding and decoding cellular information through signaling dynamics. Cell 152, 945–956 (2013).

26. Peng, L. et al. Memory hierarchy performance measurement of commercial dual-core desktop processors. J. Syst. Archit. 54, 816–828 (2008).

27. Elani, Y., Gee, A., Law, R. V. & Ces, O. Engineering multi-compartment vesicle networks. Chem. Sci. 4, 3332–3338 (2013).

28. Wauer, T. et al. Construction and Manipulation of Functional Three-Dimensional Droplet Networks. ACS Nano 8, 771–779 (2014).

29. Elani, Y., Law, R. V. & Ces, O. Vesicle-based artificial cells as chemical microreactors with spatially segregated reaction pathways. Nat. Commun. 5, 1–5 (2014).

30. Dupin, A. & Simmel, F. C. Signalling and differentiation in emulsion-based multi-compartmentalized in vitro gene circuits. Nat. Chem. 11, 32–39 (2019).

31. Pingoud, V. et al. On the Divalent Metal Ion Dependence of DNA Cleavage by Restriction Endonucleases of the EcoRI Family. J. Mol. Biol. 393, 140–160 (2009).

32. Yang, C. J., Li, J. J. & Tan, W. Using molecular beacons for sensitive fluorescence assays of the enzymatic cleavage of nucleic acids. in Methods in Molecular Biology vol. 335 71–81 (Humana Press Inc., Totowa, NJ, 2006).

33. Crisalli, P. & Kool, E. T. Multi-path quenchers: Efficient quenching of common fluorophores. Bioconjugate Chem. 22, 2345–2354 (2011).

34. Marras, S. A. E. Selection of fluorophore and quencher pairs for fluorescent nucleic acid hybridization probes. in Methods in Molecular Biology vol. 335 3–16 (Humana Press Inc., Totowa, NJ, 2006).

35. Walker, B., Kasianowicz, J., Krishnasastry, M. & Bayley, H. A pore-forming protein with a metal-actuated switch. Protein Eng. Des. Sel. 7, 655–662 (1994).

36. Braha, O. et al. Designed protein pores as components for biosensors. Chem. Biol. 4, 497–505 (1997).

37. Chang, C. yu, Niblack, B., Walker, B. & Bayley, H. A photogenerated pore-forming protein. Chem. Biol. 2, 391–400 (1995).

38. Bayley, H. Pore-Forming Proteins with Built-in Triggers and Switches. Bioorg. Chem. 23, 340–354 (1995).

39. Koo, S., Cheley, S. & Bayley, H. Redirecting Pore Assembly of Staphylococcal α-Hemolysin by Protein Engineering. ACS Cent. Sci. 5, 629–639 (2019).

40. Hammerstein, A. F., Jayasinghe, L. & Bayley, H. Subunit dimers of α-hemolysin expand the engineering toolbox for protein nanopores. J. Biol. Chem. 286, 14324–14334 (2011).

41. Dixit, S. S., Pincus, A., Guo, B. & Faris, G. W. Droplet shape analysis and permeability studies in droplet lipid bilayers. Langmuir 28, 7442–7451 (2012).

