## Supplementary Information for "A lipid-based parallel processor for chemical signals"

### Supplementary Figures

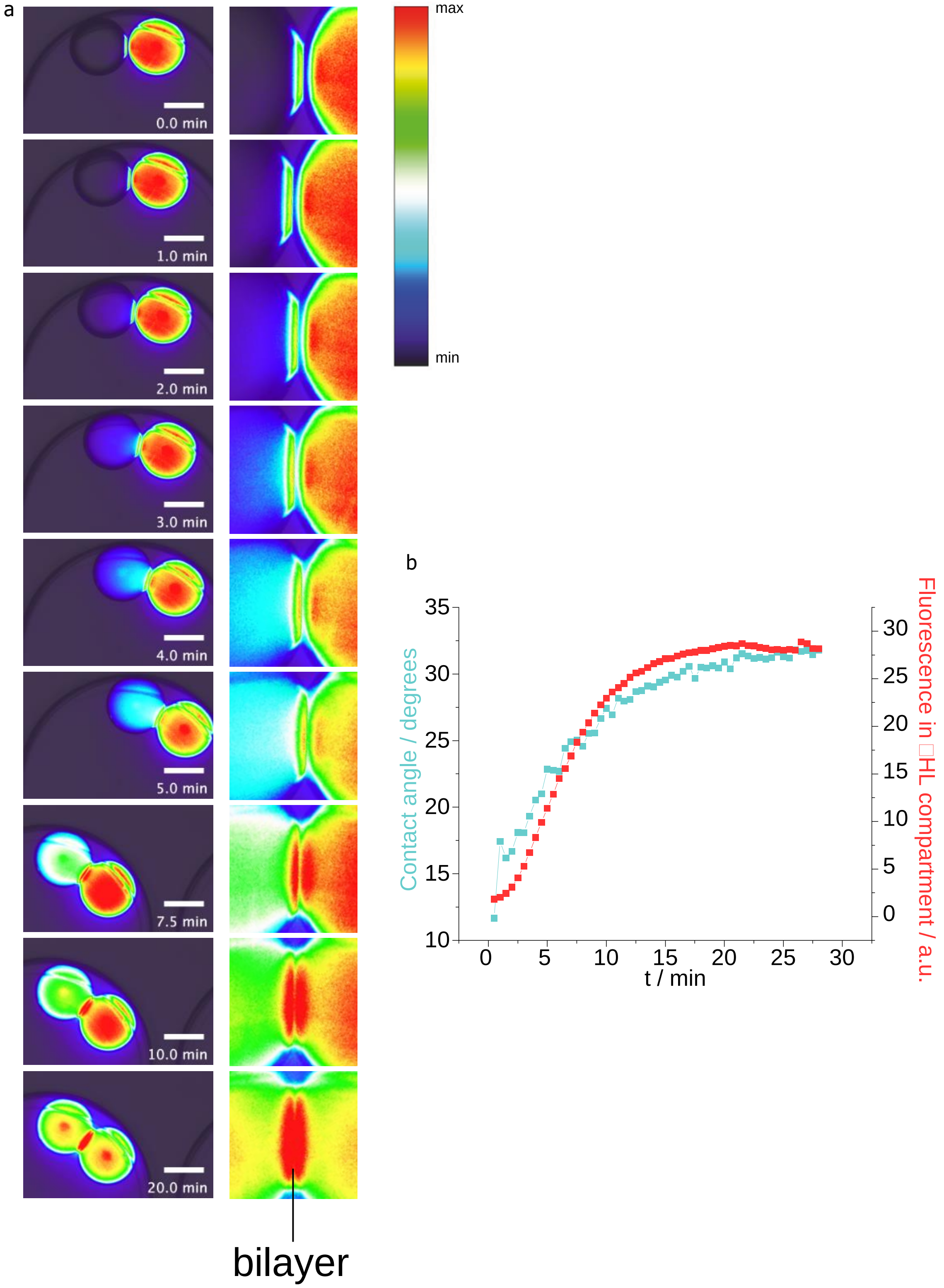


**Supplementary Figure 1. a** Composite bright-field and epifluorescence time-lapse microscopy images of molecular diffusion from a signal release compartment containing 2-NBDG into a signal transmission compartment containing purified αHL monomers, in an external lipid-oil environment, as seen in Supplementary Video 1. Magnified images of bilayers corresponding to each time point are also shown. Molecular diffusion began immediately upon contact of the compartments, before the bilayer had expanded to its final area. A plume of fluorescence was initially observed near the bilayer in the αHL-containing compartment, followed by mixing inside the compartment. Scale bars = 300 μm. **b** Plots of the contact angle between the two compartments and fluorescence in the signal transmission compartment over time. Increasing contact angle indicates increasing bilayer area. Bilayer formation began when the two compartments touched, and proceeded over several minutes. The rate of bilayer formation was high at the beginning and decreased as the bilayer approached its maximum size. The rate of molecular diffusion started low and rapidly increased as more pores formed in the bilayer. After the maximum number of pores had formed, molecular diffusion reached a constant rate. Then, as the concentration gradient of the molecules across the bilayer became smaller, molecular diffusion slowed down.

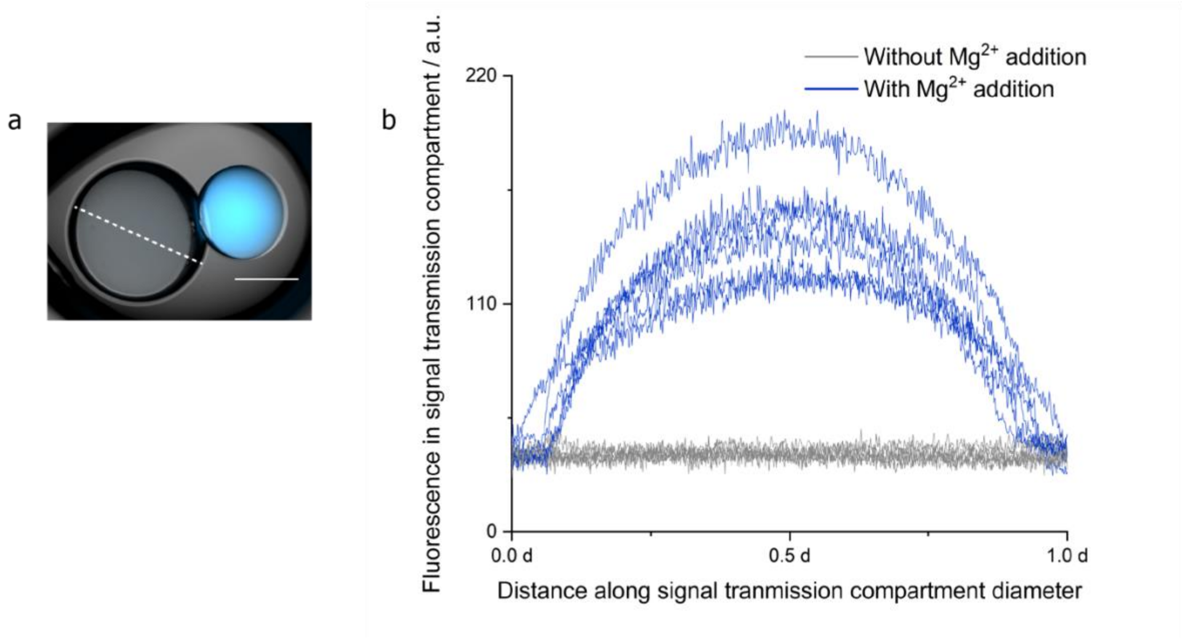


**Supplementary Figure 2.** Analysis of fluorescence in the signal transmission compartment of two-compartment processors containing an EcoRI+DNA compartment as seen in Fig. 4. **a** Fluorescence of each signal transmission compartment was analysed by drawing a line across the compartment, indicated as dashed white line. **b** Fluorescence values across signal transmission compartments, d = compartment diameter. In processors given the $Mg^{2+}$ input signal, the fluorescent product from the EcoRI reaction was observed in the signal transmission compartment ($n$ = 6). No fluorescence was observed in the signal transmission compartment if $Mg^{2+}$ was not added ($n$ = 6).

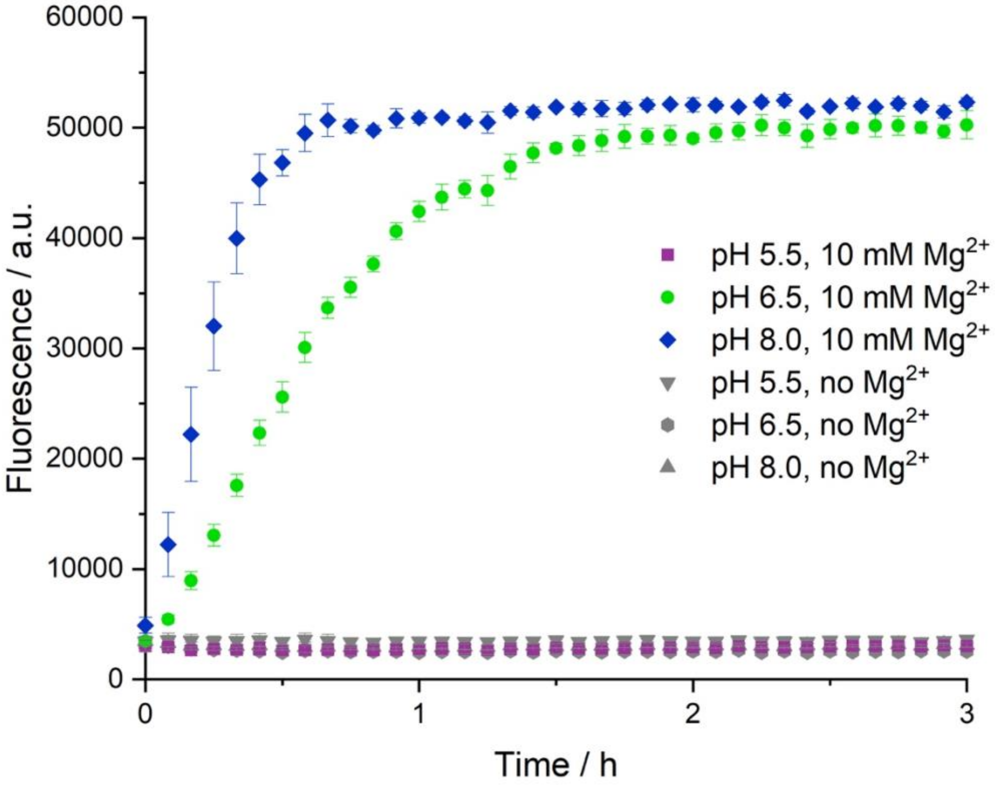


**Supplementary Figure 3.** EcoRI activity at various pH values, with or without the co-factor $Mg^{2+}$. The fluorescent product was measured. DNA cleavage by EcoRI did not proceed without $Mg^{2+}$ ($n$ = 3 for each pH condition). The EcoRI enzyme was highly active at pH 8.0 ($n$ = 3). Reduced activity was observed at pH 6.5 ($n$ = 3), whereas no activity was observed at pH 5.5 ($n$ = 3). Error bars represent standard deviation.

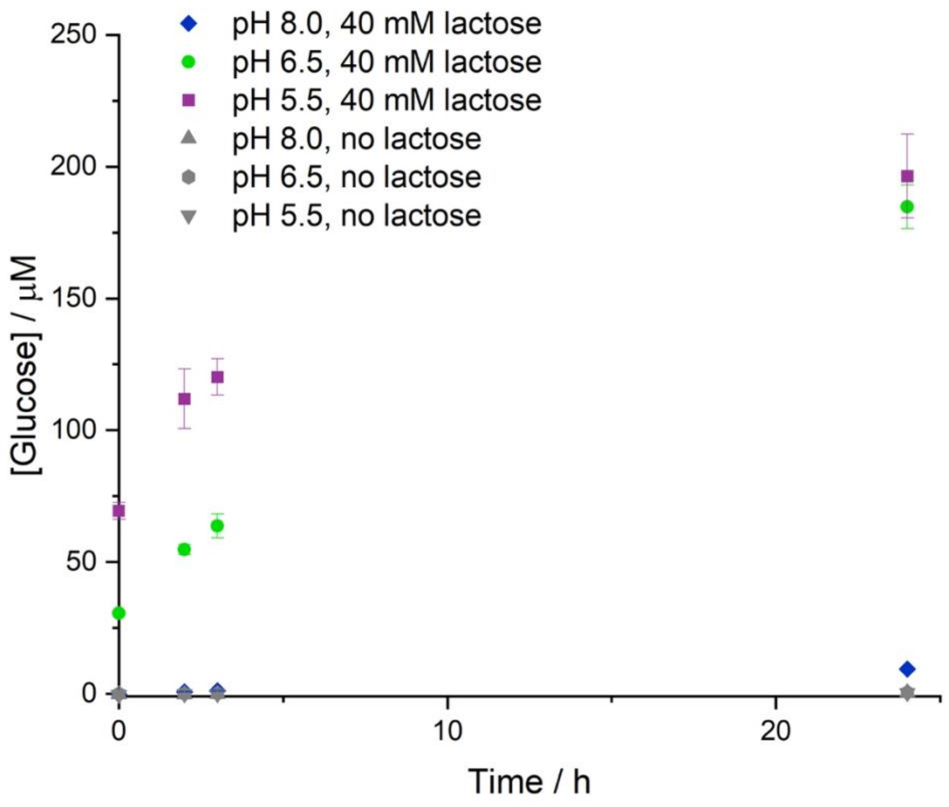


**Supplementary Figure 4.** Generation of glucose by β-galactosidase activity at various pH values, with or without the lactose substrate (*n* = 3). No glucose product was detected when the lactose input signal was not present (*n* = 3 for each pH condition). β-galactosidase was highly active at pH 5.5 (*n* = 3). Reduced activity was observed at pH 6.5 (*n* = 3), and very low activity at pH 8.0 (*n* = 3). Error bars represent standard deviation.

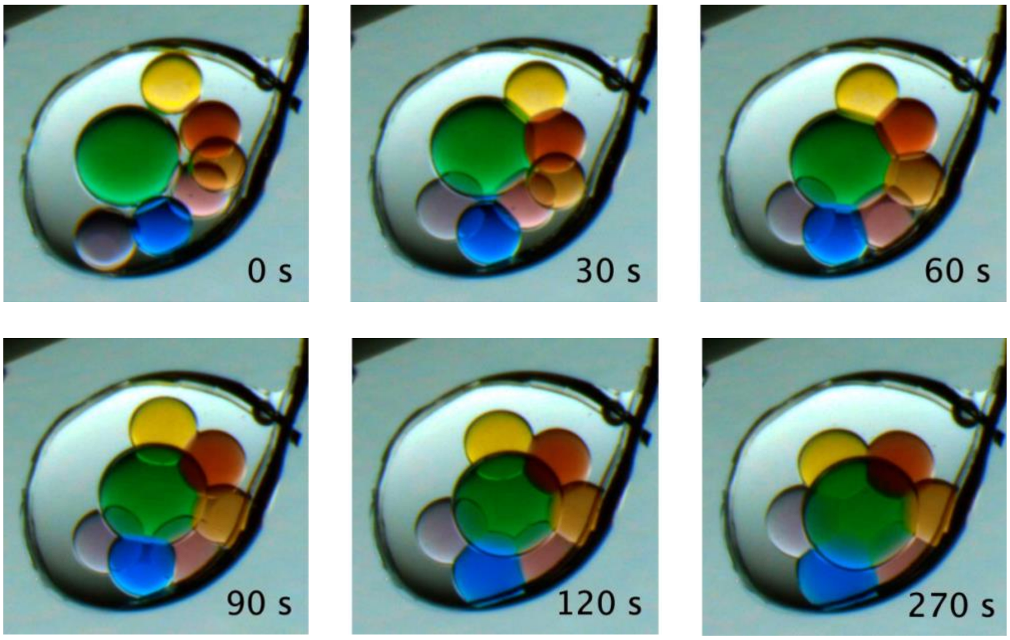


**Supplementary Figure 5.** Generation of a multi-compartment processor mimic as seen in Supplementary Video 3. A larger signal transmission compartment mimic in the middle (green dye) was connected to 6 processing compartment mimics (various colours). The final system self-assembled into the desired flower arrangement within 4.5 min. Wire diameter = 76 µm.

#### Supplementary Note

To minimise the background fluorescence of Rhod-2 ($K_d$ for $Ca^{2+} \approx 3.8$ μM) due to trace metal ions in the aqueous solutions, we included 2 μM of a non-fluorescent chelator, 1,2-bis(o-aminophenoxy)ethane-N,N,N′,N′-tetraacetic acid) (BAPTA, $K_d$ for $Ca^{2+} \approx 160$ nM) inside the sensing compartment and external aqueous environment. Unlike Rhod-2, BAPTA was not dextran-conjugated and could diffuse through the αHL pores. Over time, excess BAPTA from the external solution diffused into the sensing compartment and competed with the Rhod-2 to bind the trace ions, further reducing the fluorescence of Rhod-2.

### Descriptions of Additional Supplementary Files

**Supplementary Video 1.** Composite bright field and epifluorescence microscopy time lapse of molecular diffusion from a signal release compartment containing 2-NBDG into a signal transmission compartment containing αHL monomers, within an external lipid-oil environment, as seen in Supplementary Fig. 1. Each compartment was 410 μm in diameter. Molecular diffusion began immediately upon contact of the two compartments and proceeded simultaneously with bilayer formation. The fluorescence signal was mapped onto a spectrum colour map as indicated in Supplementary Fig. 1. Scale bars = 300 μm.

**Supplementary Video 2.** Time lapse of molecular release from a two-compartment structure, which comprised a signal release compartment (250–300 μm in diameter) containing 2-NBDG and a signal transmission compartment (500–650 μm in diameter) containing αHL monomers, within an external aqueous environment, as seen in Fig. 3b. The formation of a bilayer between the αHL compartment and the external aqueous environment was observed as a growing circle in the middle of the αHL compartment. Molecule release through the 2 bilayers was complete within 10 min.

**Supplementary Video 3.** Time lapse of the formation of a multi-compartment processor mimic with one large signal transmission compartment and six small processing compartments, as seen in Supplementary Fig. 5. Within 270 s, the compartments self-assembled into a flower arrangement with each small compartment connected to the large compartment. Wire diameter = 76 μm.
